# Taxonomic resolution of the ribosomal RNA operon in bacteria: Implications for its use with long read sequencing

**DOI:** 10.1101/626093

**Authors:** Leonardo de Oliveira Martins, Andrew J. Page, Ian G. Charles

**Affiliations:** Quadram Institute Bioscience, Norwich Research Park, Norwich, NR4 7UQ, UK.

## Abstract

Long-read sequencing technologies enable capture of the full-length of ribosomal RNA operons in a single read. Bacterial cells usually have multiple copies of this ribosomal operon; sequence variation within a species of bacterium can exceed variation between species. For uncultured organisms this may affect the overall taxonomic resolution, to genus level, of the full-length ribosomal operon.

---

Metagenomic sequencing of amplified regions of 16S, 23S ribosomal RNA genes and the ITS spacer region has become a cheap, routine and direct way to gain a high-level understanding of microbial taxonomic diversity within a sample (e.g. faeces), without the need to isolate and culture the microbes present (Quast et al. 2013; Cole et al. 2014). For example, evaluating short reads of the short hypervariable regions of the 16S gene gives family-level taxonomic resolution (Chakravorty et al. 2007). As the cost of long-read sequencing continues to fall, it is set to replace short-read sequencing, and will allow for single reads to span a whole ribosomal operon. However, the taxonomic resolution achievable using long-read sequencing of the entire length of the ribosomal operon, which encompasses the 16S and 23S genes (and usually the 5S gene), is not well explored.

As only a small number of species of bacteria are currently culturable, taxonomically classifying organisms falling into the candidate phylum radiation to species level using these methods is likely to have a higher misclassification rate.

Full-length 16S RNA can be recovered using short-read platforms (Burke and Darling 2016), and paralog copies can be discriminated, reducing chimerism and primer bias (Karst et al. 2018). The 16S-ITS-23S region of the rRNA operon has four times the variability of the 16S region alone, and can be used to classify sequences taxonomically, even at the strain level in certain cases (Benítez-Páez and Sanz 2017).

BY analysing all paralogous copies at once, we found that even when the full ribosomal RNA operon was available without chimerism or other assembly artefacts, the choice of which genomic copy to analyse affected the phylogenetic inferences; this was not only at the strain level, but at the species level ― see below. To quantify the evolutionary information from distinct rRNA segments we designed a tree silhouette score, which is a measure of how well an evolutionary tree represents taxonomically-related strains. The fact that the choice of full-length rRNA gene copy has such a strong influence reinforces the importance of capturing sufficient genetic variability to inform strain-level specificity; this cannot be properly addressed using ribosomal amplicon-based sequencing. Accounting for underlying genetic diversity has implications not only for taxonomic classification, but also for downstream analyses at the ecological or evolutionary level; in the current study we show that this has not have been fully explored previously.

Using a comprehensive data set of Gram-negative bacteria, we were able to compare the advantages and disadvantages of analysing only hypervariable regions of the 16S gene or using all three genes on the ribosomal RNA operon (16S, 23S, 5S), either independently or concatenated in all possible combinations (e.g. ‘16S5S’ represents the aligned sequences from the 16S and the 5S genes concatenated for each strain). Since there can be several copies of each operon in the same genome, we just used the longest copy in our analysis, although the results were very similar when we used a consensus of all copies (see Methods).

In cluster analysis, the silhouette score averaged over all strains is used as an indication of clustering fitness. However, we can also look at the score for each individual strain, and here we describe a simple ‘tree silhouette score’ using the patristic distance (i.e. path length between strains in the estimated phylogeny). This allowed us to quantify how well the tree retained the taxonomic classification at the species level whilst also providing evolutionary information. The distributions of silhouette scores for maximum likelihood trees estimated using i) hypervariable regions of the 16S gene, ii) entire genes independently and iii) concatenated gene sequences can be seen in Fig. 1 (top panel). We also generated a random tree with the same leaf names for comparison. All data sets (i), ii) and iii)) performed in a similar way in terms of the taxonomic resolution that could be achieved, with the exception of the 16Sv1v2 region and the 5S gene; for these two the estimated trees showed an overall poor congruence with the established taxonomic information. The 16Sv4 segment had high scores for a few strains, but it also provided a high false positive rate for other strains. By computing the proportion of samples with ‘good’ scores (i.e. above zero, which indicates correct clustering), it was apparent that longer sequences led to better taxonomic resolution overall (Supplementary Figure 1).

**Figure 1:**
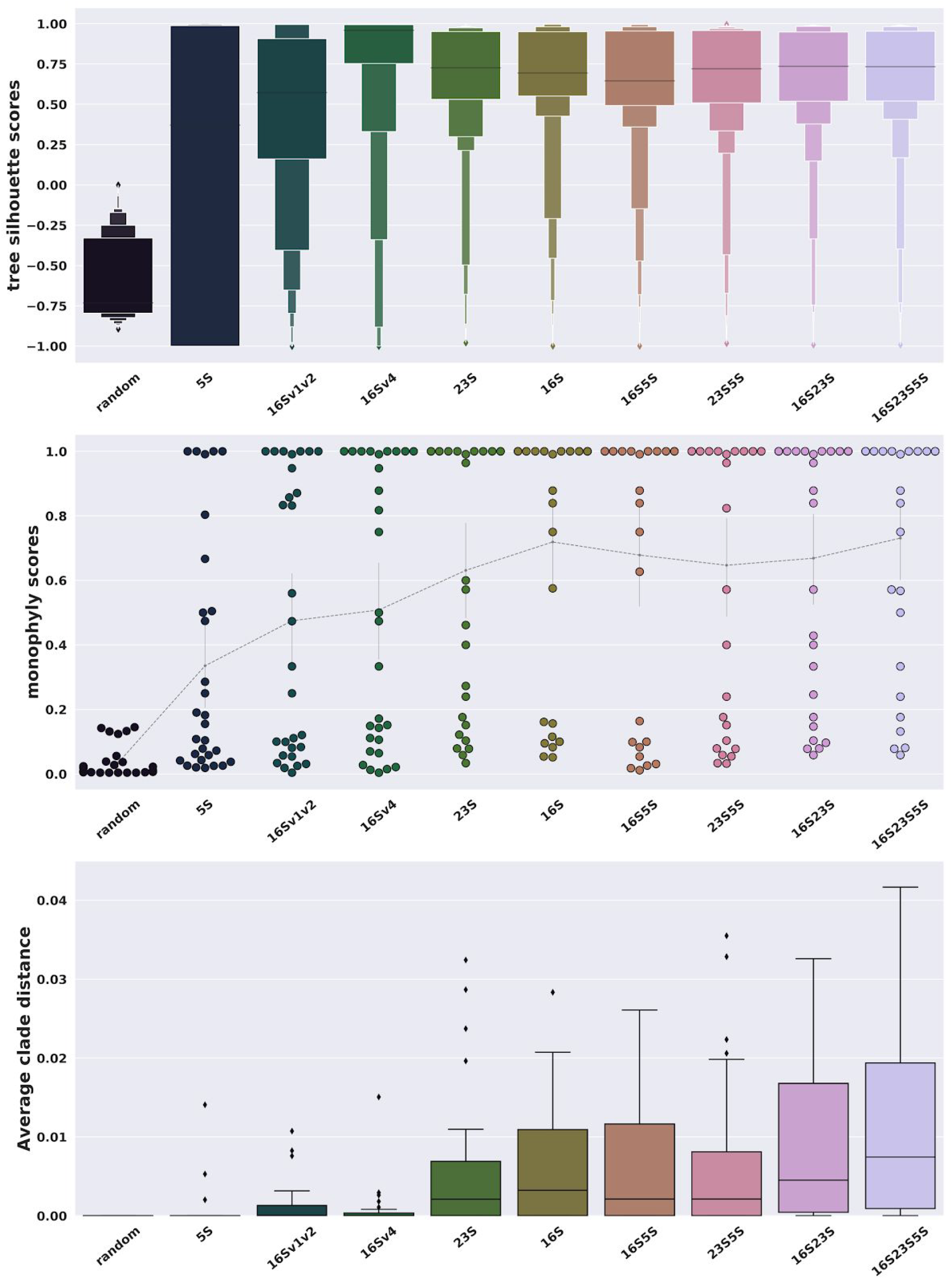
Silhouette and monophyly scores. Average branch lengths of monophyletic clades used ML trees of the longest copy of the operon. This shows the changes in phylogenetic and taxonomic resolution apparent when either: i) just a fragment of a gene (regions v1+v2 or region v4 of the 16S rRNA); ii) a whole gene; or iii) several concatenated genes are analysed. The silhouette score is at the top; it describes how close each strain is to others from the same species, compared to the closest strain from a different species. Monophyly scores are in the middle panel; they are the fraction of strains from the same species below their last common ancestor. The average patristic distance between monophyletic strains is shown at the bottom; it is the average distance between strains below the most diverse monophyletic clade of each species.

We also calculated the monophyletic status of a species directly by looking at the last common ancestor of all strains from each species, and seeing how often those ancestors also have descendants from other species. The distribution of monophyly scores for all species is shown in Figure 1 (middle panel); longer sequences resulted in phylogenetic inferences more consistent with the taxonomic classification. In particular, the concatenated data set 16S23S5S has the lowest number of species with a monophyly score lower than 0.5; such low values mean that, from the most recent common ancestor of a species, it is more common to find descendant strains from other species than from its own species. Also, for the longest concatenated data sets, the estimated evolutionary distance between samples from the same species in monophyletic clades was larger than when using smaller sequences (Figure 1, bottom). Overall this means that longer alignments can provide richer phylogenetic signals than short alignments, without compromising taxonomic resolution.

Species of bacteria often contain multiple copies of the ribosomal operon, and each has its own distinct phylogenetic diversity; intragenomic variability in the 16S gene has been studied most commonly (Větrovský and Baldrian 2013; Coenye and Vandamme 2003), but there are also studies on the 23S and 5S genes (Pei et al. 2012, 2009). When we consider the evolutionary history of all copies of the ribosomal RNA operons from a given species, one underlying assumption is that all those copies should be monophyletic, i.e. they should be closer to each other than to operons from other species. This is the justification for using just one copy, or a consensus of all operons as sufficient for phylogenetic inference and reconstruction of the correct groupings. However, this is not the case as ribosomal gene copies can be mobile between operons. Sequence variation amongst ribosomal operons from one species of bacterium can be greater than inter-species variation (Johansen et al. 2017).

The choice of operon affects the phylogeny at the strain level i.e. the inference of the closest sister taxon is dependent on the choice of the paralog. As an example, each operon-copy within a particular *S. hyicus* strain (labelled 900636345), clustered with a distinct strain of *S. simulans* (Fig. 2). The other strains of *S. hyicus* clustered together with *S. pseudintermedius* and *S. schleiferi*. This is a good example, but this behaviour is quite common; even *S. simulans* strains in this example do not form monophyletic groups. The same behaviour was observed between *S. epidermidis* and *S. warneri*, and between *S. argenteus* and *S. aureus*. Similar results were observed when applying the same analysis with *Pseudomonas*, particularly for *P. fluorescens*, *P. chlororaphis* and *P. putida*, where the location of certain samples depended on which paralog operon was chosen (data not shown). This agrees with previous studies showing the importance of accounting for the intra-individual diversity of the 16S gene (Větrovský and Baldrian 2013; Coenye and Vandamme 2003). Here we show that this issue cannot be resolved by sequencing larger regions of the genome at the same time. This behaviour is not due to stochastic errors in the inference, but to underlying biological processes such as gene duplications in the sequences analysed (with differential losses) and, to a lesser degree to lateral transfers and incomplete lineage sorting (Szöllosi et al. 2014)

**Figure 2:**
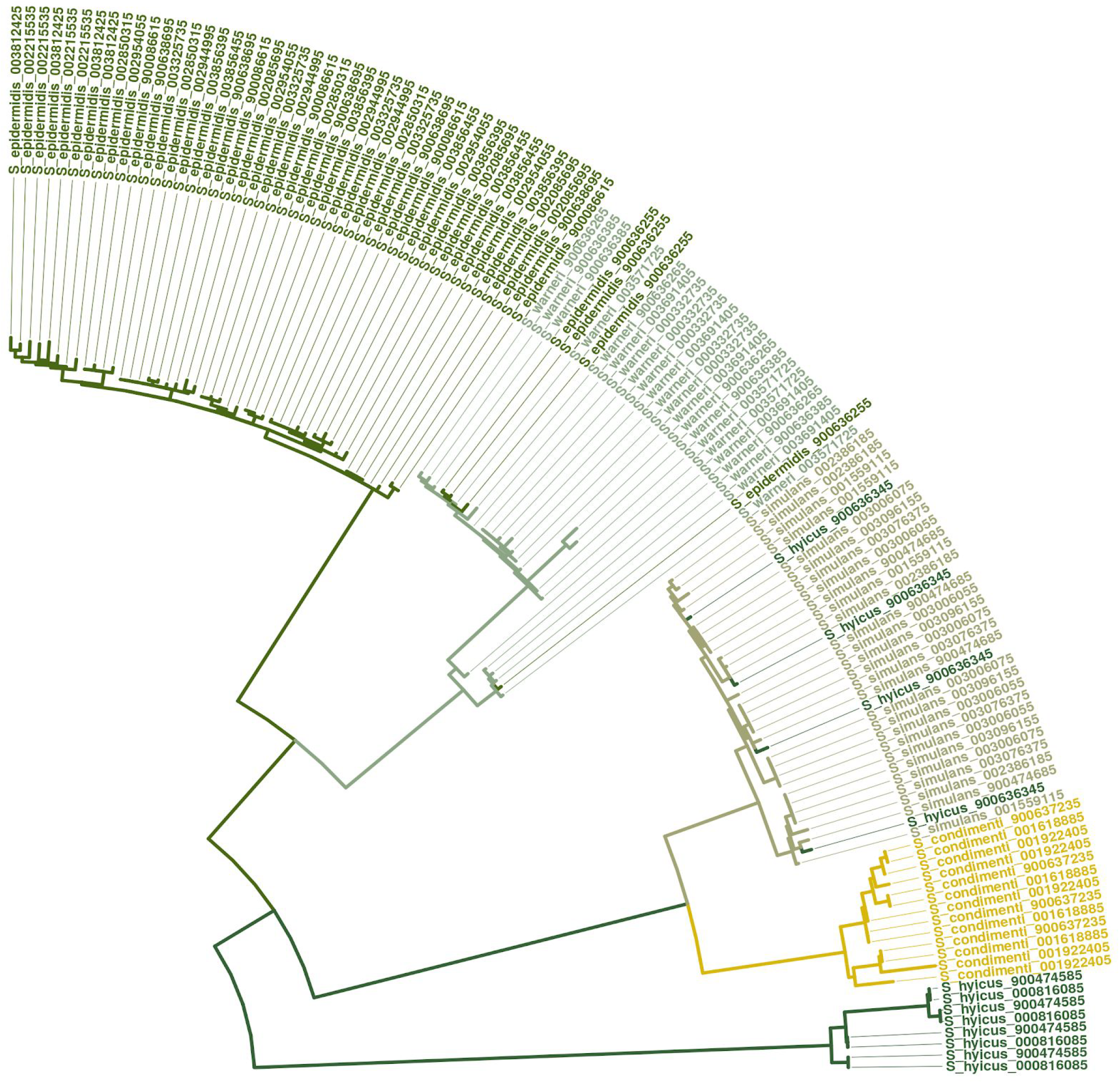
Maximum likelihood tree of full operons, including paralogs, with colours representing species. Only the region of interest is shown to emphasise the uncertain location of *S. epidermis* and *S. hyicus*: phylogeny would vary depending on the copy selected.

## Acknowledgements

LOM & AJP were supported by the Quadram Institute Bioscience BBSRC funded Core Capability Grant (project number BB/CCG1860/1). IGC was supported by the Quadram Institute Bioscience BBSRC Strategic Programme: Microbes in the Food Chain (project number BB/R012504/1).

## Methods

### Single copy data sets

For analysis of single-copy data sets, we downloaded all complete bacterial assemblies from RefSeq (accessed on 15-02-2019) of a selected set of gram-negative bacteria. We analysed all those with more than three strains from the same species, while also down-sampling over-represented ones. The selected genera were *Escherichia*, *Helicobacter*, *Leptospira*, *Neisseria*, *Pseudomonas*, and *Salmonella*. For species with more than 100 strains (namely *Escherichia coli, Salmonella enterica, Pseudomonas aeruginosa* and *Helicobacter pylori*) we assigned a probability of being chosen inversely proportional to their representativity in the database, such that we had, on average, 100 strains in the final data set. When a strain contained several ribosomal RNA operons, we chose the longest operon to provide a single sequence for phylogenetic analysis. The sequence lengths were 1539±32 for the 16S gene, 2912±89 for the 23S gene and 116±1 for the 5S gene. The related statistics for all genomes, including the number of copies per genome, can be found in the Supplementary Material.

The hypervariable regions chosen where the v1+v2 and the v4 segments of the 16S gene. They were found by assuming a 1500 bp 16S gene and then taking the first 360 bp (16Sv1v2), and the region from bases 540 to 820 (16Sv4) (Chakravorty et al. 2007); this meant that for a 16S gene of any size the first 24% of sites (360/1500) were chosen as segment v1v2 and the sites comprising the interval between 36% (540/1500) and 55% (820/1500) were assigned to segment 16Sv4. These, as well as the single genes 16S, 23S, and 5S, were aligned independently, encompassing multiple common genomic regions used for taxonomic classification. Furthermore, the multi-locus data sets like 16S+23S and others were created by concatenating the previously aligned single-locus genes.

In real-world metagenomic datasets, it is not always possible to unambiguously distinguish between distinct copies of the operon within a single strain because algorithms essentially create a chimeric or consensus sequence between all the copies. We simulated this scenario by calculating the consensus sequence between all copies, but then used the IUPAC ambiguity code rather than calling the ambiguity code N when the most frequent base cannot be acessed. This allowed us to preserve more information than would have been possible using typical methods, that mistake duplicates along the genome with polymorphism within a single copy. In terms of the tree silhouette score, this creation of consensus sequences did not affect the results when compared to the longest sequence, and are therefore not shown.

### Evaluation ― tree silhouette score and monophyly score

We implemented both tree silhouette scores and monophyly scores for this study. The silhouette score is a measure of how close a sample is to others from the same cluster (species or genera, in our case), while at the same time how far away it is from samples from other clusters. It is commonly used in cluster analysis to define the optimal number of clusters. The silhouette score of strain *i* from cluster (species) *K* is defined as:

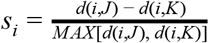

where *d* (*i*, *K*) is the average distance between strain *i* and all other strains from the same cluster (species) *K*; and *d*(*i*, *J*) is the average distance between *i* and all strains from the closest distinct cluster (species) *J ≠ K*. By definition the score *s*_*i*_ is zero if there are no other strains in the same cluster (species) *K*. In our context, the chosen distance is the patristic distance and the clusters are the species as described in the database. The patristic distance between two strains is the total path (sum of branch lengths) along the phylogenetic tree between the two leaves representing the strains. It corresponds to the cophenetic correlation coefficient in cluster analysis (Farris 1969).

The tree silhouette score as described above does not represent the phylogenetic information fully, since very short branches for sister strains (from the same species) will lead to very high scores but will have less information than one with longer terminal branches. Therefore, we also calculated silhouette scores using a simplified version of the patristic distance, sometimes called ‘path difference’, which neglects estimated branch lengths and just gives the number of internal nodes between two leaves in a tree. The fraction of strains with a positive value for this simplified score is given in Supplementary Figure S1, and estimates the fraction of strains that are closer to another strain in the same species than to one from a different species.

Besides the silhouette scores, we also implemented two statistics explicitly based on the monophyletic status of the species: the monophyly score and the best monophyletic clade score. The monophyly score of a species is the fraction of strains below its last common ancestor that are in fact from this species. In other words, for each species we find the most recent common ancestor amongst all strains from that species, and then see if there are also other species below this ancestor on the tree. The score based on the most diverse monophyletic clade is the average patristic distance between all samples below monophyletic clades. A monophyletic clade is one for which all leaves below it belong to the same species, and the average patristic distance between these leaves estimates its phylogenetic divergence. If, for a given species, we found several monophyletic clades, we chose the most diverse one, i.e. the one with the highest average distances. Higher values represent more phylogenetic information (more substitutions per site) while maintaining taxonomic resolution. In both cases we minimised spurious clades by midpoint-rerooting the trees. Note that this ‘best monophyletic clade’ is similar, but not identical to the ‘largest taxonomically consistent subtree’ (LTCS) described by Segata et al. (2013).

To have an idea of the distribution of scores in the absence of a phylogenetic signal, we generated a random tree with the same leaf names (i.e. same taxonomic information) and same tree length as the original data set. Generation of random tree branch order and lengths was achieved under a simple coalescent model using the Dendropy package (Sukumaran and Holder 2010). The same software was used for all tree manipulations.

### Operon multi-copy analysis

To study the influence of paralogy on the rRNA-based classification, we simulated the effect of long-read-based sequencing (operon multi-copy analysis) by extracting all full operons from the best represented *Staphylococcus* species in the RefSeq database. *Staphylococcus* was chosen as it is a well-studied, clinically important, pathogen. This was done by finding all annotated 16S, 23S, and 5S genes in the genome and merging all genes that were closer than 1000 bp from each other, into a single operon. For example, if the first base of a 23S gene was less than 1000 bp downstream of a 16S gene, then they would belong to the same operon, together with all sites between them. In this way we reconstructed the 16S-ITS-23S-5S operon which represents the maximum resolution achievable under real-world conditions.

Similarly, we selected all samples from *Staphylococcus* species with more than two reference genomes in the RefSeq database (accessed on 15-02-2019), while downsampling the over-represented species, *S. aureus*. Since we included all copies (average of 4.25 operons per sample), this gave rise to 167 genomes spanning 710 rRNA operon sequences. More information can be found as Supplementary Table 3. The operon lengths were between 3105 and 6186 bp (mean 5133 bp with standard deviation of 287). We then aligned all sequences with MAFFT (Katoh and Standley 2013) (automatic algorithm selection and offset of 0.3) and estimated the maximum likelihood tree with IQTREE (Nguyen et al. 2015) using the HKY+gamma evolutionary model, replicating best practice (Lees et al. 2018). The aligned sequences had 7410 columns. We repeated the same analysis with *Pseudomonas*, and all scripts necessary for reproducing these analyses are available from https://github.com/quadram-institute-bioscience/70S-resolution.

## Supplementary Tables

All supplementary tables can be found as supplementary material and at https://docs.google.com/spreadsheets/d/1XWV6N9P46UkxJo066tyhMtkpYDHeTm9OIDObc1YawM8/edit?usp=sharing

**Supplementary Table 1:** Accession numbers, species names, strain names, and summary statistics for the 758 gram-negative samples used in the single-copy analysis of the 70S rRNA sequences. The statistics shown are the number of copies, length of longest copy, and length of consensus for each of the three rRNA genes. The size of the consensus sequence is usually larger than the longest sequence since it represents an alignment of all copies (where insertions/deletions are introduced).

**Supplementary Table 2:** Accession numbers, species names, strain names and statistics for the 200 *Pseudomonas* genomes used in the operon analysis (the alignment has 960 sequences of 7803 sites in total). The statistics are the number of operons, as well as the minimum, mean, and maximum operon lengths.

**Supplementary Table 3:** Accession numbers, species names, strain names and statistics for the 167 *Staphylococcus* genomes used in the operon analysis (the alignment has 710 sequences of 7410 sites in total). The statistics are the number of operons, as well as the minimum, mean, and maximum operon lengths.

**Supplementary Table 4:** Overall statistics on sequences lengths for selected rRNA genes

## Supplementary Figure

**Supplementary Figure 1:**
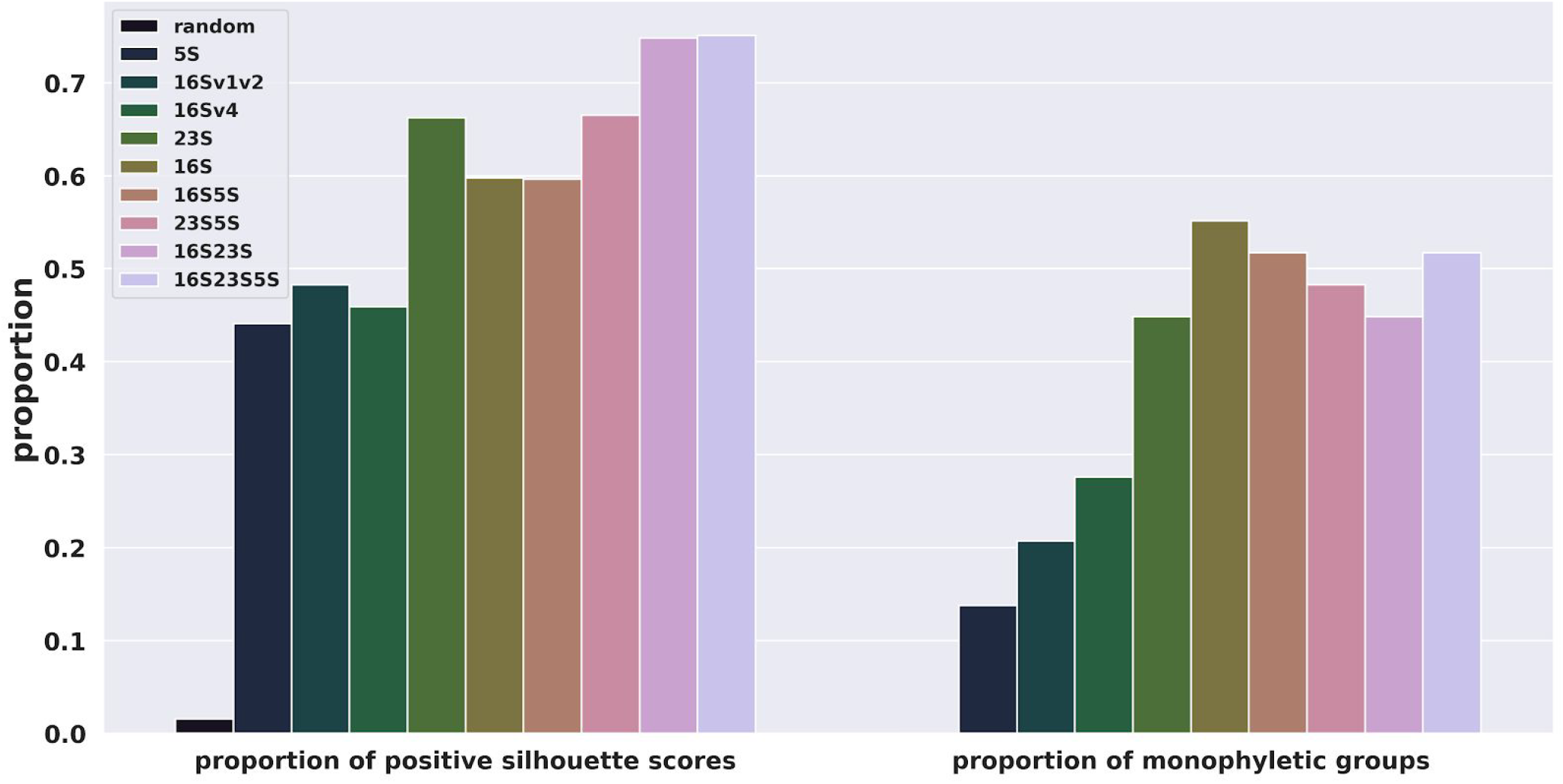
Proportion of samples with positive silhouette scores (using the simplified distance) and proportion of monophyletic species, according to maximum likelihood trees estimated from distinct combinations of rRNA sequences.

